# Cortical state fluctuations during sensory decision making

**DOI:** 10.1101/348193

**Authors:** Elina A. K. Jacobs, Nicholas A. Steinmetz, Matteo Carandini, Kenneth D. Harris

## Abstract

Neocortical activity varies between states of “synchronization” and “desynchronization”, with desynchronized states believed to occur specifically in regions engaged by the task. To disambiguate whether desynchronization is linked to task performance or engagement, we trained mice on tasks in which incorrect responses due to disengagement (neglect) differed from inaccurate task performance (incorrect choices). Using widefield calcium imaging to measure cortical state across many areas simultaneously, we found that desynchronization was correlated with engagement rather than accuracy. Consistent with this link between desynchronization and engagement, we found that rewards had a long-lasting desynchronizing effect. To determine whether engagement-related changes in cortical state depended on the sensory modality, we trained mice on visual and auditory task versions and found that desynchronization was similar in both and more pronounced in somatomotor than either sensory cortex. We conclude that variations in cortical state are predominately global and closely relate to variations in task engagement.

## INTRODUCTION

Both in sleep and in wakefulness, the cerebral cortex operates in multiple states, which can be distinguished by their degree of synchronization. In the most alert conditions, cortical activity exhibits a “desynchronized state” of relatively steady activity. As animals become less alert, the cortex enters a more “synchronized” state, characterized by low frequency (0.1-3 Hz) fluctuations in population activity that can also be detected in electroencephalogram or local field potential recordings (Buzsáki and Draguhn, 2004; Harris and Thiele, 2011). Multiple lines of evidence suggest that cortical state correlates with performance in behavioral tasks, but the precise nature of this relationship is not yet clear.

One hypothesis for the relationship between cortical state and behavioral performance is that desynchronized states enhance sensory processing. Several studies have reported that cortical representations of sensory stimuli are more faithful in desynchronized states due to lower noise correlations, even if response sizes are not always larger (Castro-Alamancos, 2004; Goard and Dan, 2009; Marguet and Harris, 2011; McGinley et al., 2015; Pinto et al., 2013; Reimer et al., 2014; Schölvinck et al., 2015; Vinck et al., 2015; Zagha et al., 2013). The relationship of cortical state to sensory-driven behavior however is not clear. While some studies have reported increased performance in desynchronized states (Beaman et al., 2017; Bennett et al., 2013; Engel et al., 2016; Pinto et al., 2013), others have reported no difference (Sachidhanandam et al., 2013), or optimal performance in intermediate states (McGinley et al., 2015).

A complicating factor in interpreting these results is that poor performance in a behavioral task does not necessarily imply impaired sensory processing. If a subject does not respond to a stimulus, this may reflect lack of motivation or engagement, rather than sensory errors. Such ambiguity cannot be resolved in “go/no go” tasks, which confound impairments in perception of a stimulus from “lapses”, when the subject may have perceived the stimulus correctly, but was not motivated to respond to it.

Here we investigated these questions by studying how cortical state correlates with behavioral performance in a 2-alternative choice task, which allowed us to distinguish perceptual errors from differences in task engagement. We trained mice on tasks requiring different sensory modalities, and imaged population activity across dorsal cortex using widefield imaging of genetically encoded calcium indicators. We found that trials when the subject responded to the stimuli were associated with more desynchronized states. However, we saw no difference in state between correct and incorrect choices, suggesting that perception was not improved in desynchronized states. Furthermore, the desynchronization associated with increased responses was global across cortex, and not restricted to the sensory region corresponding to the stimulus modality. Neither movement nor pupil size fully explained the fluctuations in brain states, suggesting the desynchronization reflected at least in part a cognitive state of engagement. Consistent with a link between desynchronization and engagement, we observed a long-lasting desynchronizing effect of reward across cortex. We conclude that in this task, cortical states reflect a general level of task engagement, rather than a specific enhancement of sensory processing.

## RESULTS

We trained mice in multiple decision-making tasks using visual and/or auditory modalities. We first present results from the purely visual task.

### The level of task engagement varies throughout a session

We trained 15 mice to perform a head-fixed visual decision-making task (Figure 1 A-B). The mice indicated whether a Gabor stimulus appeared in the left or right visual field by turning a steering wheel, and were rewarded with water for driving the stimulus to the center of the middle screen (forced choice, (Burgess et al., 2017). Some mice (8/15) were additionally trained to give a no-go response during zero-contrast trials by keeping the steering wheel still (unforced choice, (Burgess et al., 2017; Sridharan et al., 2014). Trials were classified into three groups: correct trials (turning the wheel in the direction required to receive a reward); incorrect trials (turning the wheel in the opposite direction); and neglect trials (no response before the trial timed out, despite presence of a stimulus). As described below, the primary difference in cortical states was observed between neglect trials and trials where the subject made a choice (correct or incorrect), but not between correct and incorrect trials. We will therefore group together correct and incorrect trials as “choice” trials for many analyses.

**Figure 1.**
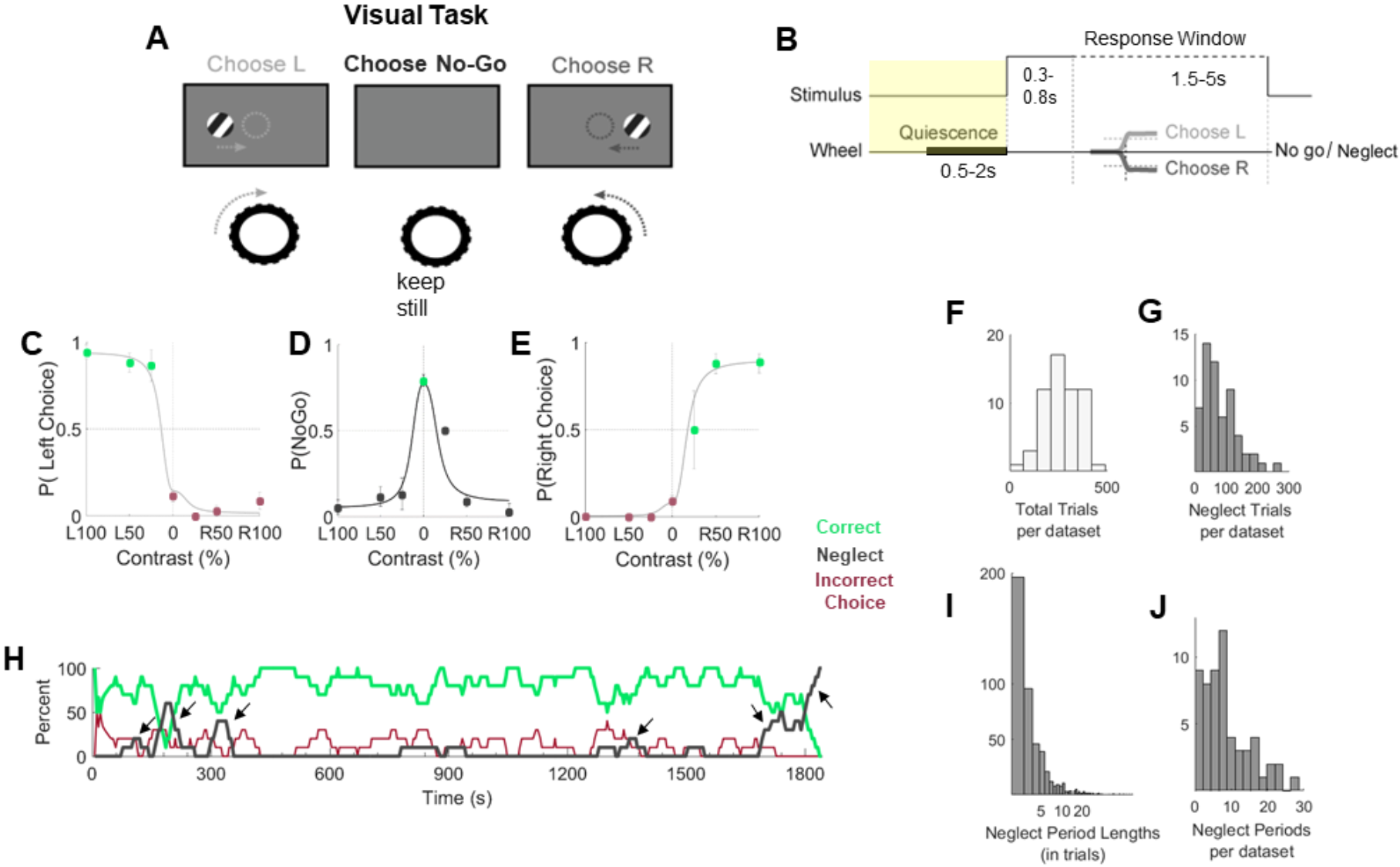
Level of engagement varies throughout a session. **A.** The visual task: a Gabor grating of varying contrast appears on the left or right visual field. The mouse must move the wheel to center the stimulus to receive a reward. The absence of a stimulus indicates a no-go trial, during which the animals are required to keep the wheel still. **B.** Timeline of the trial structure: all trials started with a baseline of 1-5 seconds (yellow highlight). Animals had to remain quiescent for 0.5-2seconds to initiate the appearance of a stimulus. After the stimulus appeared, a go cue signaled the start of the response window. If animals did not make a choice within the response window, a no-go or neglect response was recorded. **C-E.** Example psychometric curves. Error bars represent standard error of the mean. **F-G.** Number of total and neglect trials across all datasets. **H.** Engagement over time in an example dataset that contained several short neglect sequences. Green, correct trials; red, incorrect trials; black, neglect trials. Arrows point out short neglect sequences. **I-J.** Neglect period lengths and frequencies across all datasets. A-B adapted with permission from Burgess et al., 2017.

Task sessions lasted 20-60 min during which animals completed up to 400 trials and produced high quality psychometric curves (Figure 1 C-E). Nevertheless, animals occasionally provided neglect responses (Figure 1 D, G), which often came in a sequence (Figure 1 H-J) (p<0.05, t-test comparing actual and shuffled neglect sequence lengths). These neglect sequences frequently occurred in the middle of a session: the mice disengaged for a period before re-engaging with the task.

### Engagement correlates with desynchronization in visual cortex

While the mice were performing the task, we used widefield calcium imaging to record global cortical activity. We first focused our analysis on the region of primary visual cortex retinotopically aligned to the task stimuli (Figure 2 A). Although this region’s sensory responses did not consistently differ between behavioral responses (choice versus neglect) (Supplementary Figures 1 and 2), we observed a robust relationship of trial type to the power spectrum of the calcium signal during the pre-stimulus baseline period of each trial (Figure 2 B-E). Low frequency power was greater in neglect than in choice trials, with the largest differences in the 3-6Hz frequency band (Figure 2 F). A similar decrease in low frequency power was seen across sessions and animals, regardless of which mouse line was used for GCaMP expression (Figure 2 G; p<0.001, one-sample t-test on the population) and a similar relationship was seen for all genotypes examined (ANOVA, p>0.05). We therefore conclude visual cortex is more desynchronized during periods when the animal engages with the task.

**Figure 2.**
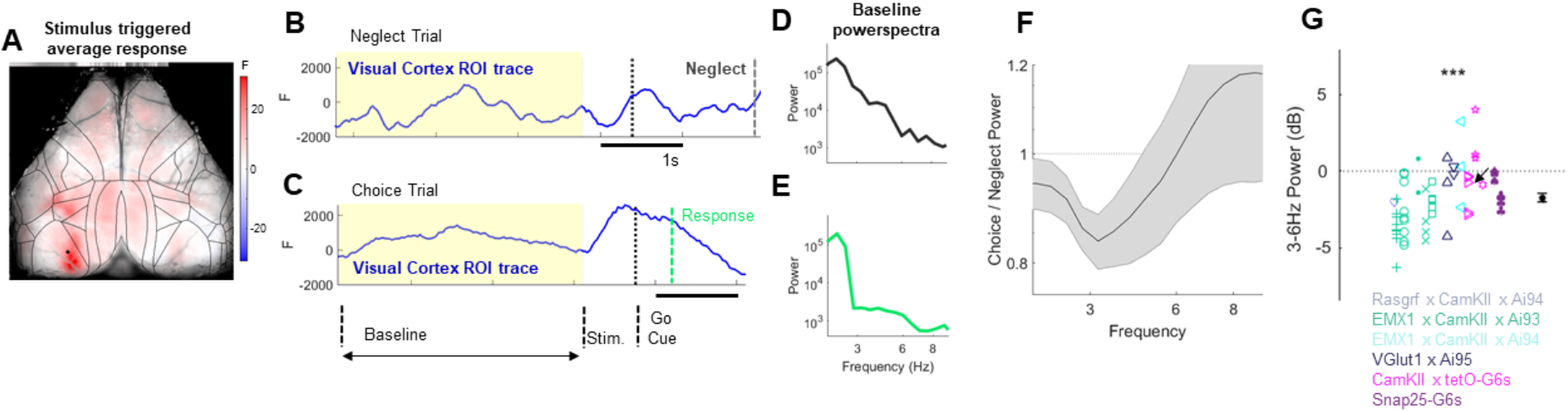
Engagement correlates with desynchronization in visual cortex. **A.** Pseudocolor representation of stimulus triggered average calcium response in 70-80ms time window after presenting high contrast visual stimuli in the right visual field. Black lines: cortical borders from Allen atlas. The dot in the left hemisphere indicates the pixel from which the traces in B originate from. **B-C.** Single trial examples from representative neglect (B) and correct choice (C) trials. Yellow background indicates baseline period, during which there was no stimulus present. Black dashed line indicates the go cue (start of the response window). Green dotted line in the choice trial indicates choice time (when the stimulus crossed the threshold in the center). Dark grey dotted line in the neglect trial indicates timeout (the animal failed to provide a choice and a neglect trial is registered). **D-E.** Power spectra computed from the baseline periods (yellow highlights in B) of the example neglect (D) and choice (E) trials. **F.** Average choice/neglect power spectra ratio in visual cortex from all experiments. Shaded areas indicate SEM. **G.** Difference in 3-6Hz power between and choice and neglect trials in visual cortex across all datasets (n = 58 experiments from 15 animals of 6 different genotypes, see Methods). Arrow indicates the result from the example dataset shown in this figure. Symbol color indicates genotype, different glyphs indicate different animals. *See also* *Figures S1* and *S2*.

### Engagement-related power differences are strongest in somatomotor cortex

We next asked whether the decrease in low frequency power during choice trials was specific to visual cortex or was a global feature of task engagement. In order to exclude possible effects of movement on brain state, we focused our analysis on the quiescent period of the baseline (Figure 3A).

**Figure 3.**
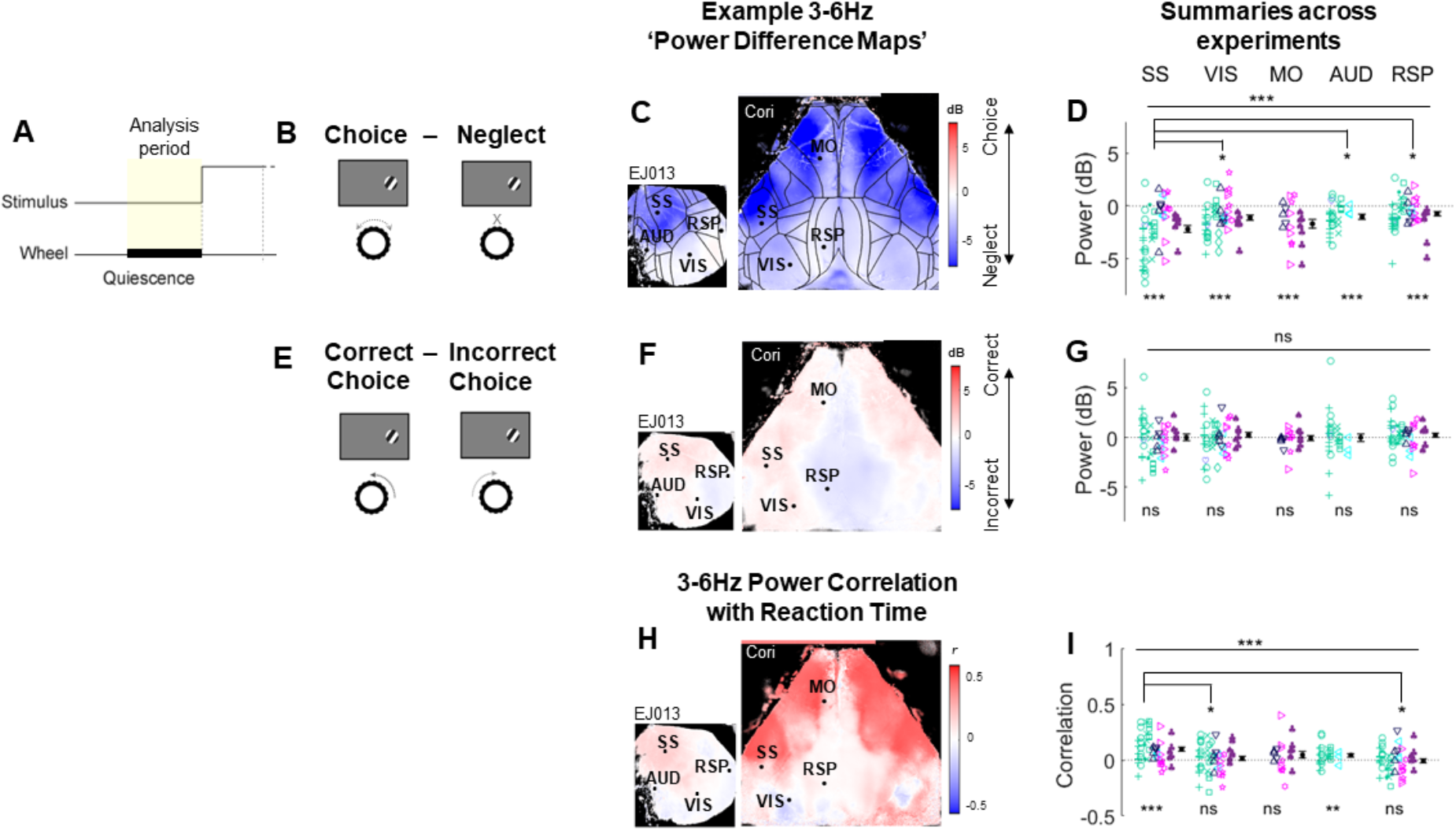
Engagement-related power differences are strongest in somatomotor cortex. **A.** Schematic indicating analysis period. **B.** Cartoon illustrating the comparison of choice and neglect power. **C**. Example maps showing the difference in 3-6Hz power between choice and neglect trials, in pseudocolor for each pixel. Blue indicates greater power in neglect trials. Left, unilateral imaging; right bilateral imaging (same dataset as previous 2 figures). Black lines: Allen atlas cortical boundaries. Dots indicate pixels that were treated as the cortical regions of interest (ROIs: Primary visual cortex = VIS, primary somatosensory cortex = SS, primary auditory cortex = AUD, retrosplenial cortex = RSP, secondary motor cortex = MO). **D.** Summary of 3-6Hz power difference between choice and neglect trials for selected ROIs across all experiments (n = 58 experiments from 15 animals). Negative values indicate stronger low frequency power preceding neglect trials. **E-G.** Similar analysis comparing correct and incorrect choices. No significant difference in power was found, for any area. **H.** Pseudocolor map showing correlation of reaction time with 3-6Hz quiescent period power in each pixel, for the same example sessions. **I.** Summary of power-reaction time correlations for all experiments. * = p < 0.05, ** = p < 0.01, *** = p < 0.001, ns=not significant. Main effect of behavioral condition (choice vs neglect, correct choice vs incorrect choice) is illustrated by the encompassing bar at the top of summary graphs and was tested using a mixed random effects ANOVA model. The effect per ROI was evaluated using student’s t-tests, and to test differences between areas we used one-way ANOVAs. *See also* *Figure S3*.

To assess brain state across as many cortical regions as possible, we employed two different imaging strategies. For some animals, we imaged the entire dorsal cortical surface bilaterally, including visual, somatosensory, motor and retrosplenial cortex; for others, we imaged the left cortical hemisphere unilaterally, to provide access to auditory cortex in addition to visual, posterior somatosensory and retrosplenial cortex.

Contrary to the hypothesis that desynchronization would be restricted to visual cortex, we observed a global decrease in 3-6 Hz power (Figure 3 C-D; p<0.001 mixed random effects ANOVA; p<0.001 one-sample t-tests per ROI). The largest effect occurred not in visual but in somatomotor cortex (p<0.001; SS vs RSP p<0.001, SS vs VIS, AUD p<0.05, SS vs MO p>0.05, one-way ANOVA). We did not observe a difference in low frequency power prior to trials where the mouse turned the wheel in the correct and incorrect direction (Figure 3 F-G) and we therefore kept them grouped together as Choice trials.

Pre-stimulus cortical state also correlated with reaction time. Choice trials with less pre-stimulus low-frequency power had faster reaction times; this correlation occurred globally (Figure 3 H-I; p < 0.001 one-sample t-test for all correlations; SS p<0.001, AUD p<0.01, VIS, MO, RSP p>0.05, one-sample t-test per ROI), with the strongest correlation in somatosensory cortex (SS vs RSP p<0.001, SS vs VIS p<0.01, SS vs MO, AUD p>0.05). We therefore conclude that cortical desynchronization correlates with task engagement: although cortical state does not predict whether the subjects will choose correctly or incorrectly, it predicts whether they will respond at all, and further predicts how quickly they will respond to the stimulus.

### Desynchronization reflects a cognitive state of engagement

By focusing our analysis on the quiescent period of the baseline, we ensured that pre-trial desynchronization was not related to pre-trial movement. However, two interpretations of our finding are possible. First, desynchronization might signal that the animals are genuinely cognitively engaged in the task; second, given the shorter reaction times during more desynchronized trials, desynchronization might simply reflect an ongoing state in which animals had an increased tendency to move.

To distinguish these alternatives, we used a task variant in which withholding a wheel movement was correct under certain task conditions, termed “2 alternative unforced choice” (2AUC; Figure 4; (Burgess et al., 2017; Sridharan et al., 2014)). In the 2AUC task, mice were rewarded for keeping the wheel still in trials where no stimulus was presented on either side. In this task, trials were therefore classified into 4 types: correct and incorrect choices (where a stimulus was present and the wheel was moved); correct no-go (where no stimulus was present and the wheel was not moved); and neglect (where a stimulus was present but the wheel was not moved). Providing a response during a no-go trial was considered an incorrect go.

**Figure 4.**
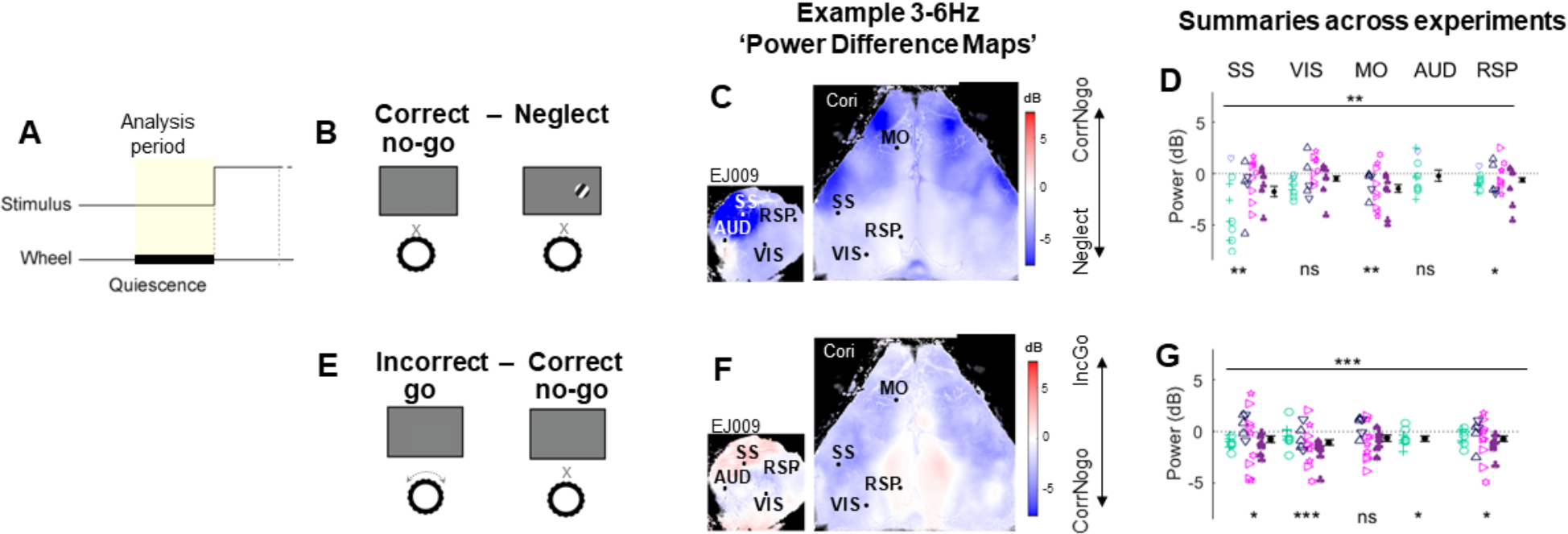
Desynchronization reflects a cognitive state of engagement. **A.** Schematic indicating analysis period. **B.** Cartoon illustrating comparison of correct no-go vs. neglect trials. **C.** Example maps showing 3-6Hz power difference; blue indicates higher power in neglect trials. **D.** Summary of 3-6Hz power difference during the quiescent period across experiments. **E-G.** Same analysis for comparison of incorrect go (incorrect choices during zero contrast trials) and correct no-go trials.

Consistent with desynchronization reflecting increased cognitive engagement rather than simply signaling upcoming movement, correct no-go trials showed significantly less 3-6 Hz power than neglect trials even though neither trial type involved movement (Figure 4 C-D; p<0.01 mixed random effects ANOVA; differences between areas not significant, p>0.05 one-way ANOVA). Nevertheless, when comparing zero contrast trials only, incorrect go trials (when the animal moved instead of keeping still for a reward) showed less 3-6Hz power than correct no-go trials (Figure 4 F-G, p<0.001 mixed random effects ANOVA). These results suggest that desynchronization reflects a difference in task engagement, that is associated with an increased likelihood of movement, which however can be successfully suppressed when required.

### Variations in cortical synchronization are not fully explained by pupil size

Pupil diameter correlates with arousal and mental effort in humans, and with cortical state in rodents (de Gee et al., 2014; Kahneman and Beatty, 1966; McGinley et al., 2015; Reimer et al., 2014). We therefore asked how pupil diameter related to engagement and cortical state in our experiments, and whether the correlation between engagement and cortical state could be explained by a common effect of pupil size.

As expected, pupil size correlated negatively with low frequency power during the baseline period: the smaller the pupil, the greater the low frequency power (Figure 5 A-C). Nevertheless, pupil size did not fully explain the state-engagement correlation: even after accounting for the effect of pupil size, there was significantly more low frequency power in neglect trials (Figure 5 A-C; p<0.05, ANCOVA). A consistent main effect of behavioral condition on low frequency power was present in all ROIs after accounting for pupil size (Figure 5 D-E; p<0.001 mixed random effects ANOVA; p<0.001 one-sample t-tests per ROI). Again, the effect in somatosensory cortex was significantly stronger than in visual, auditory and retrosplenial but not secondary motor cortex (p<0.01; SS vs RSP p<0.01, SS vs VIS, AUD p<0.05, one-way ANOVA). Similarly, in the 2AUC task, during trials with similar pupil sizes, there remained a significant difference in low frequency power between correct no-go and neglect trials (Supplementary Figure 4; p<0.001 mixed random effects ANOVA; SS,VIS, RSP p<0.001, MO p<0.01, AUD p>0.05, one-sample t-tests per ROI).

**Figure 5.**
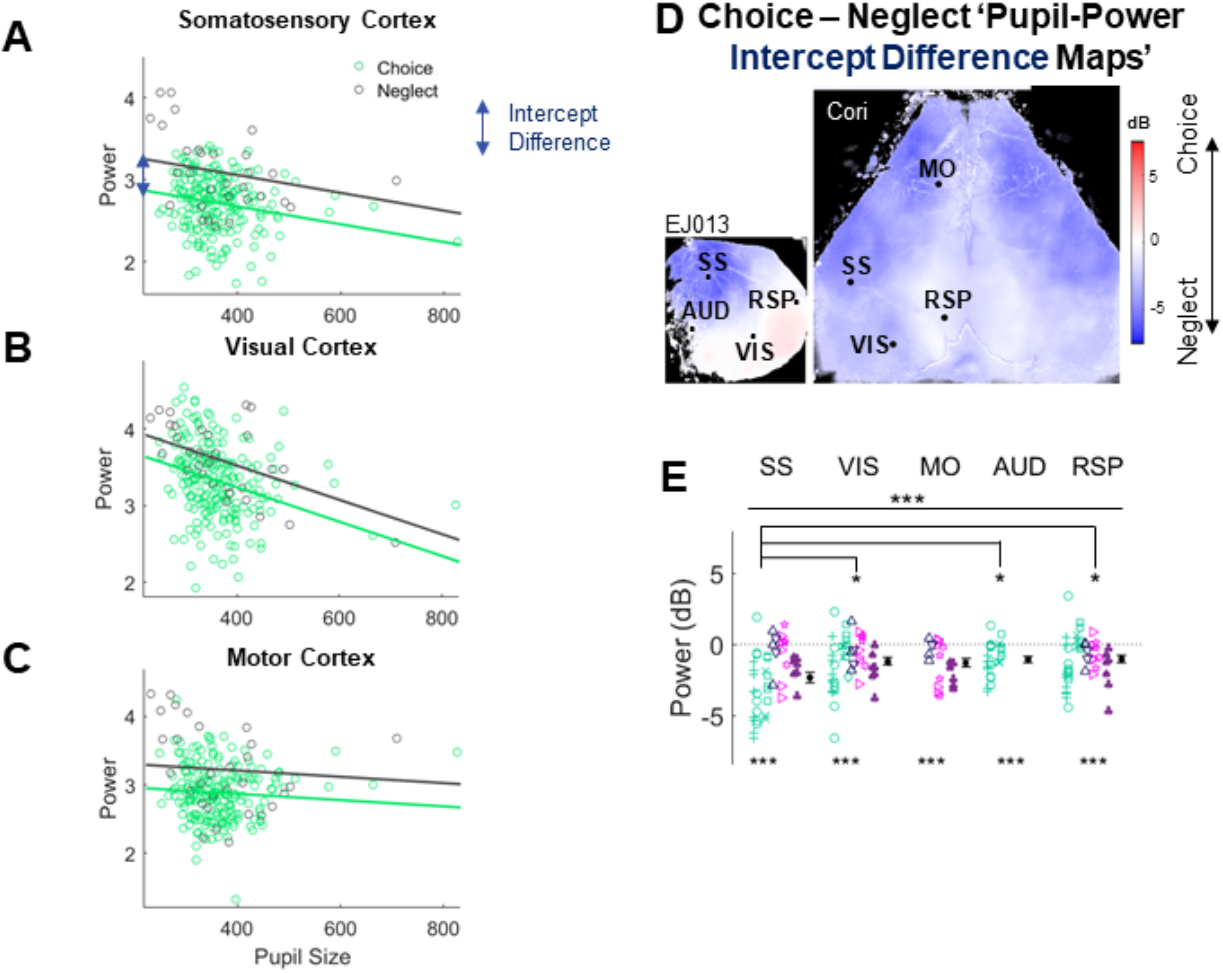
Variations in cortical synchronization are not fully explained by variations in pupil size. **A.** Relationship between pupil size, behavioral condition, and 3-6Hz power in somatosensory cortex of an example experiment. Each dot represents a trial, colored according to outcome: choice (green) or neglect (dark grey). The lines represent fits from an ANCOVA model, which captured the main effect of behavioral condition as the difference in intercept between the choice and neglect fits (blue arrow). **B-C.** Similar analysis for visual and motor cortex ROIs. **D.** Pseudocolor maps showing intercept difference for each pixel, for the same example sessions shown in previous figures. Blue indicates significantly higher power in neglect trials, after accounting for the common effect of pupil size. **E.** Summary of intercept differences across experiments in selected ROIs (n = 48 experiments from 12 animals). *See also* *Figure S4*.

### Reward is followed by prolonged desynchronization

As described above, low-frequency power did not differ between the baseline periods prior to correct or incorrect trials (Figure 3 F-G). Surprisingly however, cortical state differed in the baseline periods that followed correct and incorrect choices. As both correct and incorrect trials suggest an engaged state and were preceded by a wheel turn, but only the correct trial was rewarded, this suggests that reward was itself the factor driving the difference in cortical state.

To exclude the possibility that the physical act of reward consumption itself was affecting cortical state, we restricted our analysis to the quiescent period of the following trial (Figure 6 A & D) when the animals were no longer moving the steering wheel and by which time the animals had finished licking (as measured by a thin-film piezo sensor attached to the lick spout) in 98% of trials (6,148/6,250, data not shown). 3-6Hz power was lower in the quiescent period following correct compared to incorrect trials (Figure 6 B-C; p<0.05 mixed random effects ANOVA; SS, MO p<0.01, VIS p < 0.05, AUD, RSP p>0.05 one-sample t-tests per ROI). Similarly, low frequency power was lower following correct no-go trials (no turn followed by reward) than following incorrect trials (turn followed by no reward) (Figure 6 E-F; p<0.05 mixed random effects ANOVA; VIS p<0.05, SS, MO, AUD, RSP p>0.05, one-sample t-tests per ROI). Both these trial types were in an engaged state, and again only the rewarded trial type led to subsequent desynchronization. These results therefore suggest that reward has a desynchronizing effect on brain state that persists several seconds after reward consumption has finished.

**Figure 6.**
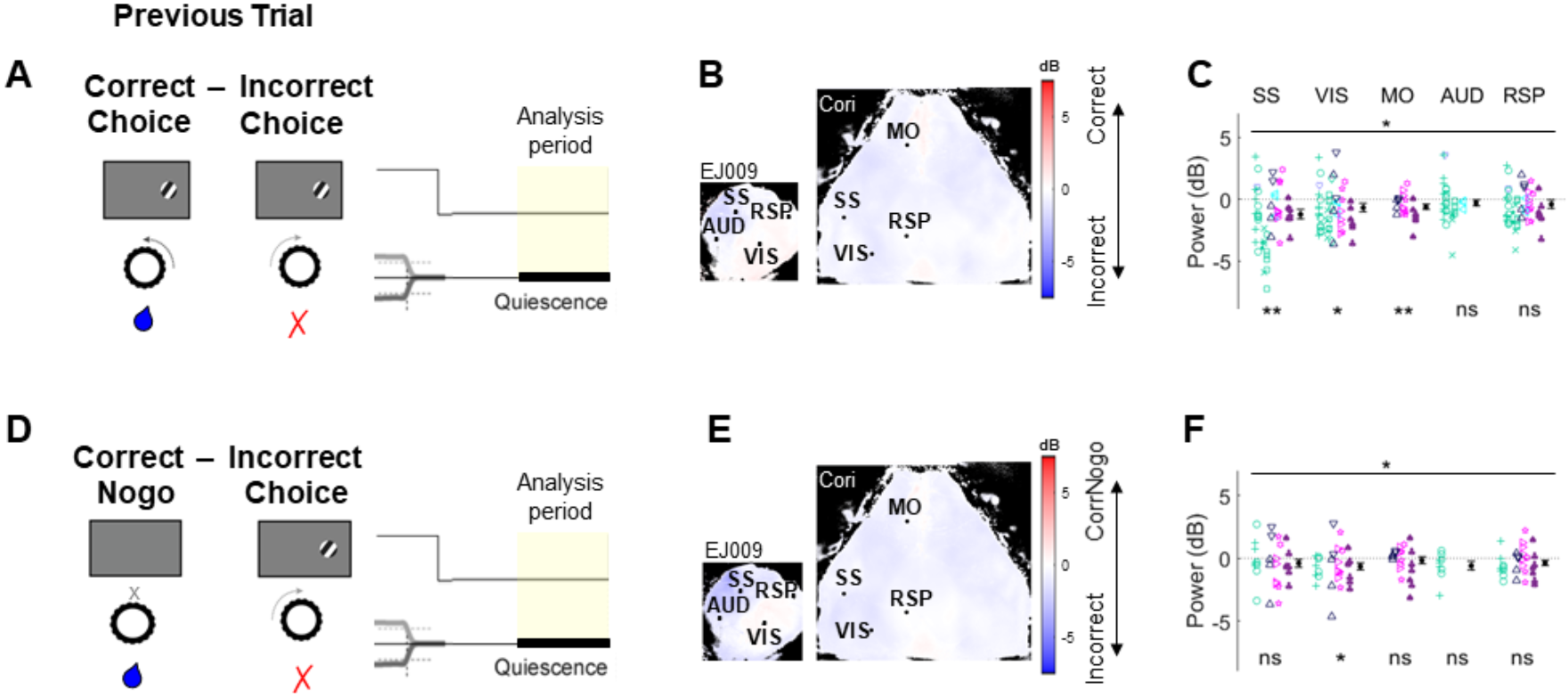
Reward is followed by prolonged desynchronization. **A.** Illustration of analysis of quiescent periods following correct and incorrect choice trials. **B.** Pseudocolor representation of 3-6Hz power difference for each pixel; blue represents lower power following correct choice. **C.** Summary across experiments for selected ROIs (n = 57 experiments from 14 animals). **D-F.** Similar analysis for comparison of correct no-go and incorrect choice trials (n = 30 experiments from 9 animals).

### Engagement-related cortical state changes are independent of sensory modality

Our results showed that task-related desynchronization was a global effect: all cortical regions desynchronized, with the strongest desynchronization seen in somatomotor rather than visual cortex. We next asked whether engaging auditory cortex by training animals in an auditory task might cause a bigger difference in auditory cortical state during task engagement. In this task, the animals were sitting in front of an isoluminant grey screen (the same screen as in the visual task), but no visual stimuli were present. Instead, the mice were presented with auditory stimuli consisting of trains of high or low frequency tones. Turning the steering wheel changed the sound frequency of the tone trains, and the mice were rewarded for bringing the stimulus frequency to a central target tone (Figure 7; Supplementary Figure 5).

**Figure 7.**
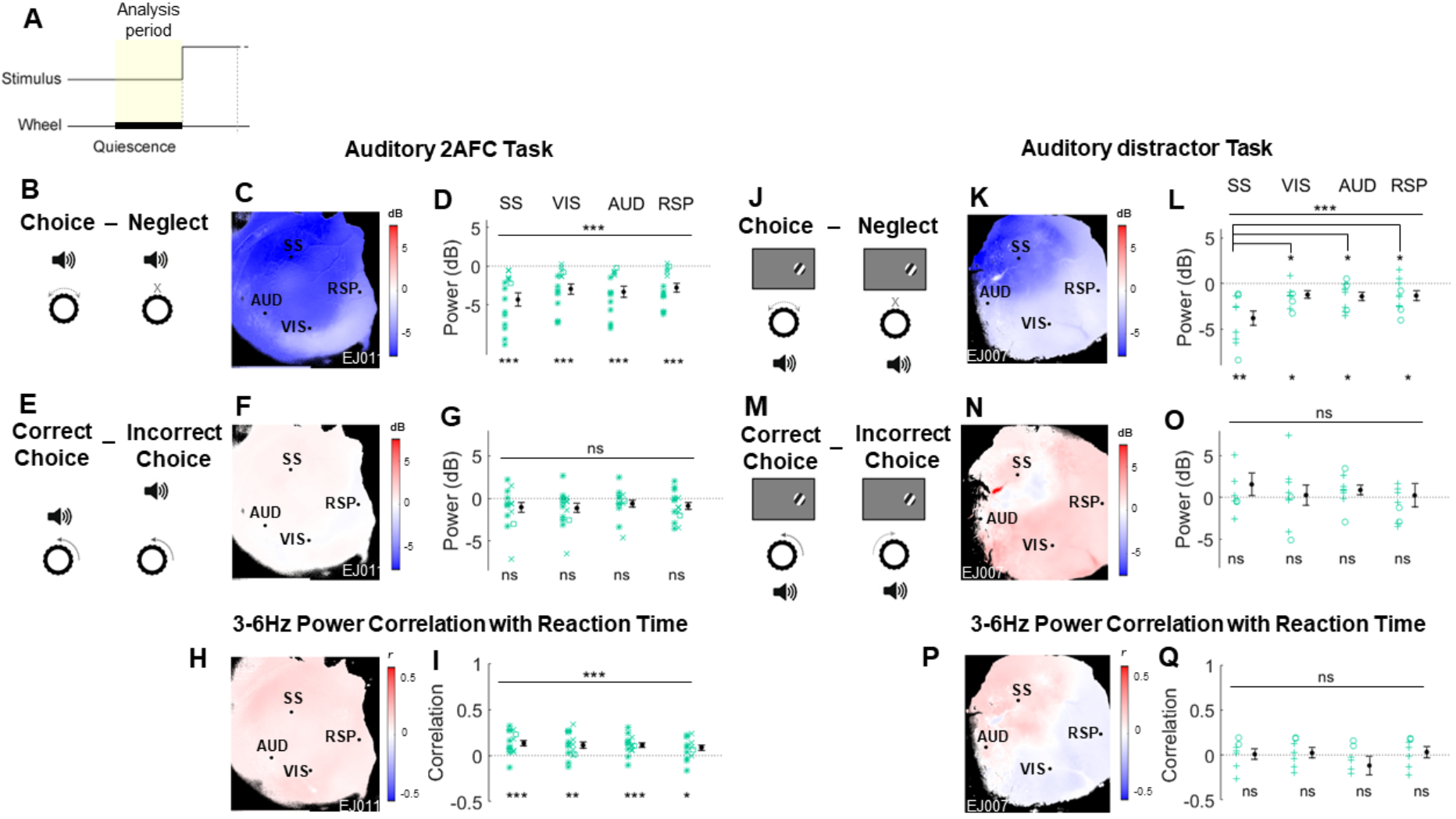
Engagement-related cortical state changes are independent of sensory modality. **A.** Schematic indicating analysis period. Trial structure is identical to the visual task, only the stimuli are different. **B.** Comparison of choice and neglect trials in the auditory 2AFC task. **C.** Pseudocolor map showing 3-6Hz power difference for each pixel; blue indicates higher power on neglect trials. **D.** Summary of 3-6Hz power difference between choice and neglect trials for selected ROIs across all experiments in the auditory task (n = 15 experiments from 3 animals). **E-G.** Comparison of correct incorrect trials in auditory 2AFC task. **H-I.** Correlation of low frequency power and reaction time in auditory 2AFC task. **J-Q.** Similar analyses for the auditory distractor task. (n = 10 experiments from 2 animals). *See also* *Figures S5*, *S6 and S7*.

In the auditory 2AFC task, desynchronization was also seen over the entire imaged window when comparing choice to neglect trials (Figure 7 C-D; p<0.001 mixed random effects ANOVA; p<0.001 one-sample t-tests per ROI). As before, this effect was more related to engagement rather than performance, as there was no significant difference in state prior to correct and incorrect trials (Figure 7 F-G). Similarly, low frequency power was significantly correlated with reaction time across the cortex (Figure 7 H-I; p<0.001 one-sample t-tests).

We also presented a subset of mice with an auditory distractor task, in which mice were presented with both visual and auditory tone train stimuli that changed in tonal frequency as the wheel was turned, but the contingency of the auditory stimulus changed between blocks, and the animals performed this task as if it was a visual task only. Mice showed similar performance on this task as in the purely visual task (Supplementary Figure 6) suggesting they disregarded the auditory stimuli.

In the auditory distractor task, brain state changes closely mirrored those in the visual and auditory tasks: we saw a global decrease in low frequency power during task engagement (Figure 7 K-L; p<0.001 mixed random effects ANOVA; SS p<0.01, VIS, AUD, RSP p<0.05 one-sample t-tests per ROI). Desynchronization was strongest in somatosensory cortex (p<0.01; SS vs VIS, AUD, RSP p<0.05, one-way ANOVA), and there were no differences in brain state prior to correct and incorrect trials (Figure 7 N-O). The correlation of low frequency power with reaction time did not reach significance (Figure 7 P-Q; one-sample t-tests), but we observed the same relationship with pupil size in the auditory and auditory distractor tasks (Supplementary Figure 7), and found the same effect of reward in the auditory task, although this did not reach significance for the auditory distractor task (Supplementary Figure 7).

## DISCUSSION

We trained mice on several discrimination tasks using visual and auditory sensory modalities. In all tasks, we found that fluctuations in engagement throughout a session correlated with cortical state. Consistent with many prior studies (Busse et al., 2017; McGinley et al., 2015; Pinto et al., 2013; Speed et al., 2018), we found that discrimination of visual or auditory stimuli desynchronizes the corresponding cortical region. However, the desynchronization was global rather than restricted to the sensory region corresponding to the stimulus modality, with the biggest effect in somatomotor cortex for all tasks.

These results seem to disagree with previous work in primate visual cortex, which showed a reduction in correlated fluctuations in parts of visual cortex corresponding to attended locations (Beaman et al., 2017; Cohen and Maunsell, 2009; Engel et al., 2016; Fries et al., 2001; Harris and Thiele, 2011; Mitchell et al., 2009). We suggest three, nonexclusive, possible explanations for this difference. First, it may reflect a difference between species. Second, while these previous studies compared different sub-regions of a single cortical area, our study compared cortical areas corresponding to different sensory modalities. Finally, it is possible that the different task demands, and correspondingly different required strategies for solving them, in those studies resulted in different cortical state changes. For example, local desynchronization may only occur when selective attention must be deployed to particular stimuli, which was not the case in our experiments. The effects observed in our experiments might have reflected arousal and engagement, which may activate more global state mechanisms, rather than selective attention, which may have more local effects.

Despite the correct choice being indicated by either visual or auditory stimuli, the region whose state showed the strongest correlation with performance was somatomotor cortex. The region where this effect was strongest corresponds to barrel cortex, which was unanticipated given that there was no overt need to use the whisker system in any of our tasks. There was no difference in overall whisker motion energy between engaged (choice) and disengaged (neglect) trials (Supplementary Figure 8), however there may have been subtle differences in whisker trajectory we could not measure with our videographic methods. It is unlikely that the mice were using their whiskers to feel the steering wheel, as they were raised sufficiently high above the steering wheel that their whiskers did not touch it during resting. Yet, given that whisking is such an ethologically important part of a mouse’s behavior (Crapse and Sommer, 2008), they may have used their whiskers in our tasks even though there was no direct need. Alternatively, it is possible that cognitive engagement, while influencing activity across cortex, influences activity most strongly in somatomotor and particularly barrel cortex for reasons that remain unknown. Lastly, it is possible that the desynchronization in somatomotor cortex reflects a state of readiness to provide a response to the sensory stimuli in order to obtain a reward. This may even be independent of the sensory processing, which could be occurring in parallel to a motor preparation.

It has been suggested that the desynchronized state improves the signal-to-noise ratio of the neural code by reducing correlated fluctuations in neural activity, thereby allowing more accurate decisions (Cohen and Maunsell, 2009; Mitchell et al., 2009). It is possible that a contrast detection task like ours does not depend as critically on cortical state as orientation change detection (Cohen and Maunsell, 2009) or spatial tracking (Mitchell et al., 2009). Nevertheless, the fact that cortical state is similar prior to correct and incorrect choices in our task challenges the hypothesis that low correlation is necessary for accurate sensory representation. From a theoretical perspective, noise correlations do not always impair sensory processing, but only do so if their population-level structure matches that of signal correlations (Averbeck et al., 2006), which may not be the case in mouse visual cortex (Stringer et al., 2018). Although desynchronization did not correlate with correct choices, it was associated with an increased speed and probability of movement, whether in a correct or incorrect direction. We therefore suggest that desynchronization represents a state of generic task engagement or motivation rather than of improved sensory processing: in behavioral states associated with cortical desynchronization, animals respond to stimuli more readily, leading to faster reaction times and decreased neglect. Importantly, animals can still successfully suppress movements in this state, as correct no-go trials are also characterized by desynchronization. Furthermore, desynchronization was seen immediately following reward delivery, consistent with increased motivation at these times.

We therefore suggest that global cortical desynchronization, rather than a state specialized for accurate sensory processing, is a state that is associated with producing rapid and coordinated behavioral responses to sensory stimuli of any modality. The movements the mouse must make in this task involve large regions of the body; making them rapidly may require widespread desynchronization of the entire cortical surface, particularly somatomotor cortex.

## EXPERIMENTAL PROCEDURES

All experiments were conducted according to the UK Animals Scientific Procedures Act (1986), under personal and project licenses released by the Home Office following appropriate ethics review.

### Animals

All animals were on a normal daylight cycle (8am - 8 pm), and co-housed whenever possible.

The mice came from a variety of genotypes, and in main-text plots (Figures 1-6) the mouse line is indicated by symbol colour, while glyph shapes represent individual mice. Animals were offspring of double or triple transgenic crosses (males: n = 7, females: n = 9), expressing either GCamp6f or GCamp6s in cortical excitatory neurons under the following drivers (color code indicated to the right):

- Ai93; Emx1-Cre; Camk2a-tTa (n = 7, Emerald green)
- Ai94; Emx1-Cre; Camk2a-tTa (n = 1, Cyan)
- Ai94; Rasgrf-Cre; Camk2a-tTa (n = 1, Lavender)
- Ai95; VGlut1-Cre (n = 2, Navy)
- tetO-G6s; Camk2a-tTa (n = 3, Magenta)
- Snap25-G6s (n = 2, Purple)

Our main results held for all genotypes, including lines which can exhibit interictal activity (Steinmetz et al., 2017).

### Surgery

Mice underwent surgery at the age of 8-10 weeks. They were anesthetized with 2% isoflurane (Merial) in oxygen, body temperature was kept at 37°C, and analgesia was provided by subcutaneous injection of Rimadyl (1ml/0.1kg, Pfizer). The eyes were protected with ophthalmic gel (Viscotears Liquid Gel, Alcon).

In unilaterally imaged animals, the temporalis muscle was detached unilaterally to expose auditory cortex on the left hemisphere. The skull was thinned above visual, auditory and posterior somatosensory cortex using a scalpel until the external table and diploe of the bone were removed. A metal head-plate with a circular opening above posterior cortex was fixed to the cranium with dental cement (Sun Medical, Moriyama, Shiga Japan), and a 8mm coverslip was then secured above the thinned skull using UV cement (Norland Optical Adhesives #81, Norland Products Inc., Cranbury, NJ USA) with a LED UV Curing System (CS2010, Thorlabs Ltd, Ely UK).

In bilaterally imaged animals, the skull was left intact and a clear skull cap implantation following the method of Steinmetz et al. (2017) was used. A light-isolation cone was 3D-printed (Ultimaker 2+, Ultimaker B.V., The Netherlands), implanted surrounding the frontal and parietal bones and attached to the skull with cyanoacrylate (VetBond; World Precision Instruments, Sarasota, FL USA). Gaps between the cone and skull were filled with L-type radiopaque polymer (Super-Bond C&B, Sun Medical, Moriyama, Shiga Japan). The exposed skull was covered with thin layers of UV cement, and a metal headplate was attached to the skull over the interparietal bone with Super-Bond polymer.

### Behavioral tasks

Mice were trained in one of several variants of a two-alternative choice task (Burgess et al., 2017). Behavioral training started 1-2 weeks postsurgery, and all animals were handled for habituation prior to head-fixation and training on the tasks. Mice were trained to sit head-fixed in front of an LCD monitor (refresh rate 60Hz), at the bottom of which a MF1 speaker (TDT, Alachua, FL USA) was placed for auditory or auditory distractor experiments. In all but 10 experiments, the monitors were covered with Fresnel lenses to make intensity spatially uniform. The paws of the mice were resting on a steering wheel, which the animals could turn to provide a response in the tasks.

In the basic visual two-alternative forced choice (2AFC) task, a visual Gabor stimulus of varying contrasts appeared randomly in the left or the right visual field. The mouse could move the stimulus on the screen by turning the steering wheel, and was rewarded with water for moving the stimulus to a central location within a response window (1.5-5 s). Incorrect choices (i.e. wheel turns in the wrong direction) or neglect responses (i.e. failure to respond within the allowed time window) resulted in a time-out, which in some mice (10/16) was also signaled white a white noise burst. (The noise burst was dropped in later experiments as it was not necessary for good performance.)

A subset of mice were trained on a visual two-alternative unforced choice (2AUC) version of the task, which contained zero contrast trials for which the animals were required to keep still during the response window in order to receive a reward (Burgess et al., 2017; Sridharan et al., 2014). In some of these mice, stimuli could be presented on both sides, and the animals were rewarded for moving the higher contrast stimulus to the center, or at random if the contrasts were equal.

In the auditory 2AFC task, low or high frequency tone trains (8 or 15 kHz, respectively) were presented from the speaker directly in front of the mice, and the movement of the wheel was coupled to changes in the tonal frequency of the tone pips. The aim of the task was to bring the tone frequency to the mid-frequency (11 kHz), which was also presented as a go-cue.

The auditory distractor task was identical to the visual 2AFC task, but with irrelevant auditory stimuli presented simultaneously with the visual stimuli. The auditory stimuli consisted of the same auditory tones as in the auditory task, which also changed in frequency as the wheel was turned. However, low and high frequency tones were inconsistently associated with visual stimuli in different sessions, such that in a given session, low was paired with left and high with right, or vice-versa. Even though the auditory stimuli could have provided information about the stimulus within a given session, the animals did not use this to perform the task; when presented with the auditory stimuli alone, they performed at chance level (data not shown). Mice trained on the auditory distractor task had not previously learned the auditory 2AFC task.

To enable analysis of cortical state prior to trial onset, all trials started with a pre-trial baseline of 1-5 seconds. For the stimulus to appear, the animals had to remain quiescent (keep the wheel still) for 0.5-2 seconds; early movement lead to a delay in stimulus appearance. Some animals were also trained to keep still for 0.3-0.8 seconds when the stimulus appeared and wait for a go cue to give their response. In the visual tasks, the go cue consisted of either a tone or a visual Gabor stimulus at the center of the screen. The modality of the cue did not affect the behavior or the results presented, therefore these tasks were analyzed together. In the auditory and auditory distractor tasks, the go cue consisted of a tone (consisting of the target frequency in the auditory task).

Psychometric curves were generated with the same generalized linear model as in Burgess et al., 2017 (see equations 1, 2, and 3).

### Widefield imaging

To correct for hemodynamic artefacts, we used alternate-frame illumination (Ma et al., 2016). GCaMP6 fluorescence was excited with a blue LED (470nm; LEX2-B, Brain Vision or Cairn OptoLED, P1110/002/000), while on alternate frames a green or violet LED was used to measure a calcium-independent hemodynamic signal. Imaging was performed at acquisition rates of 35-50 Hz per colour, 10-19ms exposures, with 2×2 or 4×4 binning using a PCO Edge 5.5 CMOS camera and a macroscope (Scimedia THT-FLSP) with 1.0x condenser lens (Leica 10450028) and 0.63x objective lens (Leica 10450027).

Imaging was conducted at two rigs with similar set-ups. At the first set-up, the excitation light was diverted to the brain via a dichroic mirror (FF506-Di03, Semrock) and passed through a bandpass filter (FF01-482/35-25, Semrock). The green light for capturing the hemodynamic signal was provided by a ring-illuminator containing 5-6 miniLEDs (528nm; Thorlabs LED528EHP), driven with a LEDD1B driver, that was fixed around the objective. The fluorescence emitted by the brain passed through a dichroic mirror (FF593-Di03, Semrock) and an emission filter (FF01-543/50-25, Semrock).

At the second set-up, the excitation light passed through an excitation filter (Semrock FF01-466/40-25), a dichroic (425nm; Chroma T425lpxr), and 3mm-core optical fiber (Cairn P135/015/003), then reflected off another dichroic (495nm; Semrock FF495-Di03-50×70) to the brain. To capture the hemodynamic signal, the light was passed through a violet excitation filter (405nm, Chroma ET405-20x) on every other frame. Light from the brain passed through a second dichroic and emission filter (Edmunds 525/50-55 (86-963)) to the camera.

### Dimensionality reduction

Widefield movie data were compressed and denoised using the singular value decomposition (SVD). All analyses were conducted directly on the SVD-transformed data, allowing much faster execution times that would be required to process the full-pixel movies. Code for such analyses is freely available at https://github.com/cortexlab/widefield.

To compress the data, first, the 3D stack was reshaped into a 2D matrix *S* of dimensions *p* × *t*, where *p* is the number of pixels and *t* is the number of time points. Then, we performed SVD of *S*:

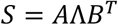

The physiological spatiotemporal dynamics of the data were fully captured with the top 500 singular values, with lower components encoding only noise. Therefore all the presented analyzes were performed using the top 500 singular values. Each pixel was expressed as a linear combination of the first 500 temporal components of Λ*B*^*T*^, which we called V; weighted by the corresponding spatial matrix U, consisting of the first 500 spatial components of *A*.

### Hemodynamic correction

Hemodynamic correction was performed by subtracting a multiple of the calcium-independent signal from the GCaMP signal. The multiple used was allowed to vary between pixels; to estimate this multiple, both GCamp and hemodynamic signals were first linearly de-trended and high-pass filtered above 0.01Hz, and then bandpass filtered in the frequency range corresponding to the heart beat (9-13Hz), where hemodynamic artifacts are strongest. The optimal multiple was estimated by linear regression; pixel-wise multiplication and subtraction was performed in the SVD domain to allow faster analysis. The code for this method can be found at https://github.com/cortexlab/widefield/blob/master/core/HemoCorrectLocal.m.

Hemodynamic correction was performed on all data except for data from 3 early animals that were imaged using blue illumination only. However, because later analyses indicated that our spectral analysis results were not affected by hemodynamic artefacts, the data from these 3 animals was also included in the paper.

### Eye tracking

Neural recordings were paired with eye tracking recordings in all but 10 datasets. One of the eyes (usually the eye contralateral to the imaged hemisphere in unilaterally imaged recordings) was illuminated with an infrared LED (SLS-0208A, Mightex; driven with LEDD1B, Thorlabs), and recorded using a 446 camera with an infrared filter and a zoom lens (Thorlabs MVL7000). The videos were recorded with MATLAB’s Image Acquisition Toolbox (MathWorks). Pupil size and position were computed following the methods from Burgess et al. (2017). All obtained pupil traces were further processed following the methods of Reimer et al. (2016).

### Behavioral measurements

Trials were divided into three classes. They were classified as “choice trials” if the animal provided a choice (correct or incorrect) within the response window. They were classified as “neglect trials” if the animal failed to provide a response within the response window, when a response was required to obtain a reward. They were considered “correct no-go” trials when the animal correctly withheld a response throughout the response window. The % correct, incorrect and timeout in Figure 1 was computed using a sliding window over 10 trials. Reaction times in Figure 3 were defined as the interval of time between go-cue onset and response time. Since reaction time varied between stimuli, we computed the average reaction time per stimulus, subtracted this average per trial per contrast, and used the obtained residuals for computing the Pearson correlation with power.

The baseline period was defined as the inter-trial interval (ITI) preceding the stimulus onset at each trial. Quiescent periods were defined as the end of the ITI during which no movement was detected. Trials with ITIs or quiescent periods of less than 0.7s were excluded from analysis.

### Stimulus-triggered responses

Trials containing 50% or higher contrast on the right visual field were averaged and baseline subtracted at time 0 of stimulus onset. The map in Figure 2A consists of the frame at t = 70-80ms post stimulus onset subtracted by the previous frame. This method was chosen as it revealed the cleanest stimulus response due to the slow dynamics of GCaMP6s.

### Power maps

As explained under ‘Dimensionality reduction’, for a given pixel *n*, the fluorescence over time was represented by 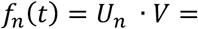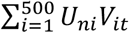, where *U*_*n*_ is the row within *U* corresponding to pixel *n*. Therefore, the Fourier transform of *f*_*n*_(*t*) was calculated as

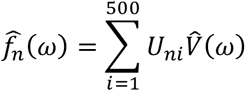

where ^ denotes the Fourier transform. To compute the power in the frequency band of interest 3-6Hz, we calculated 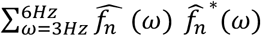. The power spectrum for all pixels could thus be efficiently computed using matrix multiplication, at least an order of magnitude faster than without the SVD compression. Finally, this was reshaped into a 2-dimensional ‘Power map’, *P*(*x,y*), where *x* and *y* are spatial dimensions.

We computed these power maps during the ITI or quiescent period for each trial separately, then computed the average power for choice and neglect conditions:

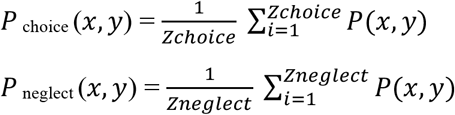

The power difference maps were then computed as follows:

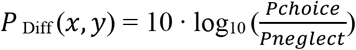

The multiplication by 10 is applied to turn the power ratios into units in decibels.

The same principle applied for computing power difference maps between correct and incorrect trials, and so on.

In the power difference maps shown in the figures, pixels with average power (over time) below the 20^th^ percentile were set to black. This procedure effectively masks pixels outside the brain.

### ROI selection

Outlines of visual and auditory cortices were identified in each mouse by sensory stimulation using sweeping bars (Kalatsky and Stryker, 2003) for visual cortex, and repeated pure tone pips of different frequencies for auditory cortex.

The region of interest (ROI) in visual cortex (VIS) was chosen as the center of the stimulus response to contralateral stimuli within the visual task. The ROI in auditory cortex (AUD) was based on the auditory cortex maps obtained by passive stimulation. The responses to different frequencies were averaged and the ROI was selected from the area with the highest mean response and which was responsive to the frequencies used within the auditory task.

The position of somatosensory cortex was estimated stereotaxically, and confirmed functionally by imaging activity during whisking and movement. The ROI in somatosensory cortex (SS) was chosen from within the area that was estimated to be the barrel cortex.

The ROI in retrosplenial cortex (RSP) was estimated stereotaxically and always chosen from posterior RSP as this was the visible part of RSP in the unilateral imaging experiments. The secondary motor cortex ROI (MO) was estimated stereotaxically.

The individual patterns of vasculature in each mouse were used as additional guidance to place the ROIs as consistently as possible across experiments from the same individuals. ROIs in the bilateral imaging experiments were chosen from the left hemisphere to be consistent with the unilateral imaging experiments.

The power for each ROI was estimated as the ‘power map’ value at a single pixel in the ROI center; in practice however, this averages a signal from a slightly larger region due to the spatial smoothing resulting from the SVD representation.

### Statistics

Unless otherwise specified, all statistical tests were carried out with a paired t-test to test the null hypothesis that the power differences or reaction time correlations across experiments come from populations with mean zero. In addition, one-way ANOVAs were employed to test the null hypothesis that the power differences did not differ between the different cortical regions of interest. Multiple comparisons were adjusted using Tukey’s honest significant difference criterion. To assess whether there was a main effect of behavior condition, a random mixed effects ANOVA model was used, in which experimental session was set as a random effect.

In figure 5, data were analyzed with one-way analysis of covariance models (ANOCOVA) fitting separate but parallel lines to the data per behavioral condition. Multiple comparisons were adjusted using Tukey’s honest significant difference criterion.

## AUTHOR CONTRIBUTIONS

E.A.K.J., M.C. and K.D.H. designed the project. E.A.K.J. and N.A.S. conducted the experiments, E.A.K.J. and K.D.H. analyzed the results, E.A.K.J. and K.D.H. wrote the manuscript, N.A.S. and M.C. reviewed and edited the manuscript.

## ACKNOWLEDGEMENTS

We thank Andrew J. Peters for help with data collection, Michael Okun for contributing code for the data analysis, Philip Coen, Laura Funnell and Miles Wells for help with animal training, and Charu Reddy for animal husbandry.

This work was supported by a Wellcome Trust PhD Studentship to E.A.K.J. (103364/Z/13/Z), a Wellcome Trust Investigator Award (205093/Z/16/Z), grants from the European Research Council (694401) and the Simons Foundation (325512), and postdoctoral fellowships from the Human Frontier Sciences Program (LT001071/2015-L) and from the European Union’s Horizon 2020 research and innovation programme under the Marie Sklodowska-Curie (grant agreement No 656528). M.C. holds the GlaxoSmithKline / Fight for Sight Chair in Visual Neuroscience.

## SUPPLEMENTARY FIGURES

**Supplementary Figure 1.**
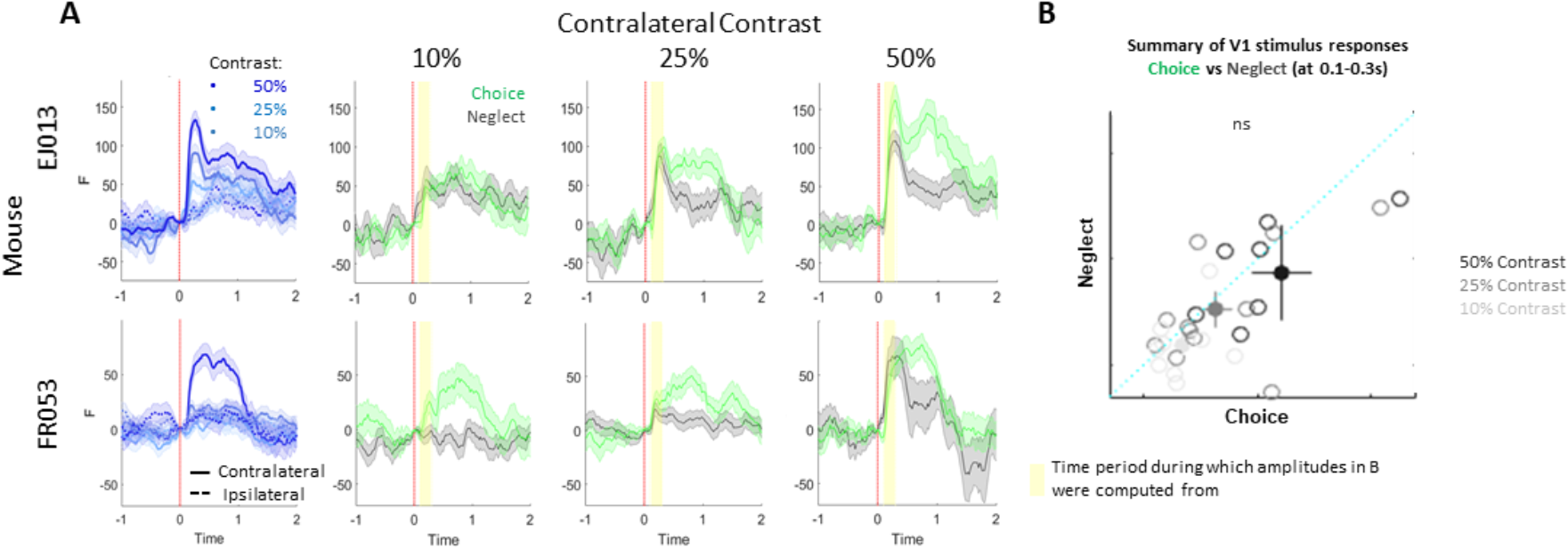
Example V1 stimulus responses from two mice. Related to Figure 2. **A.** Left-most column: Stimulus responses by contrast. Other columns: contralateral contrast responses for choice and neglect conditions. Shaded areas indicated SEM, yellow background indicates period of time period during which the stimulus response amplitudes shown in B were computed. **B.** Summary of contralateral stimulus response amplitudes across experiments. Each circle corresponds to one dataset, color denotes contrast, filled circles denote average with SEM.

**Supplementary Figure 2.**
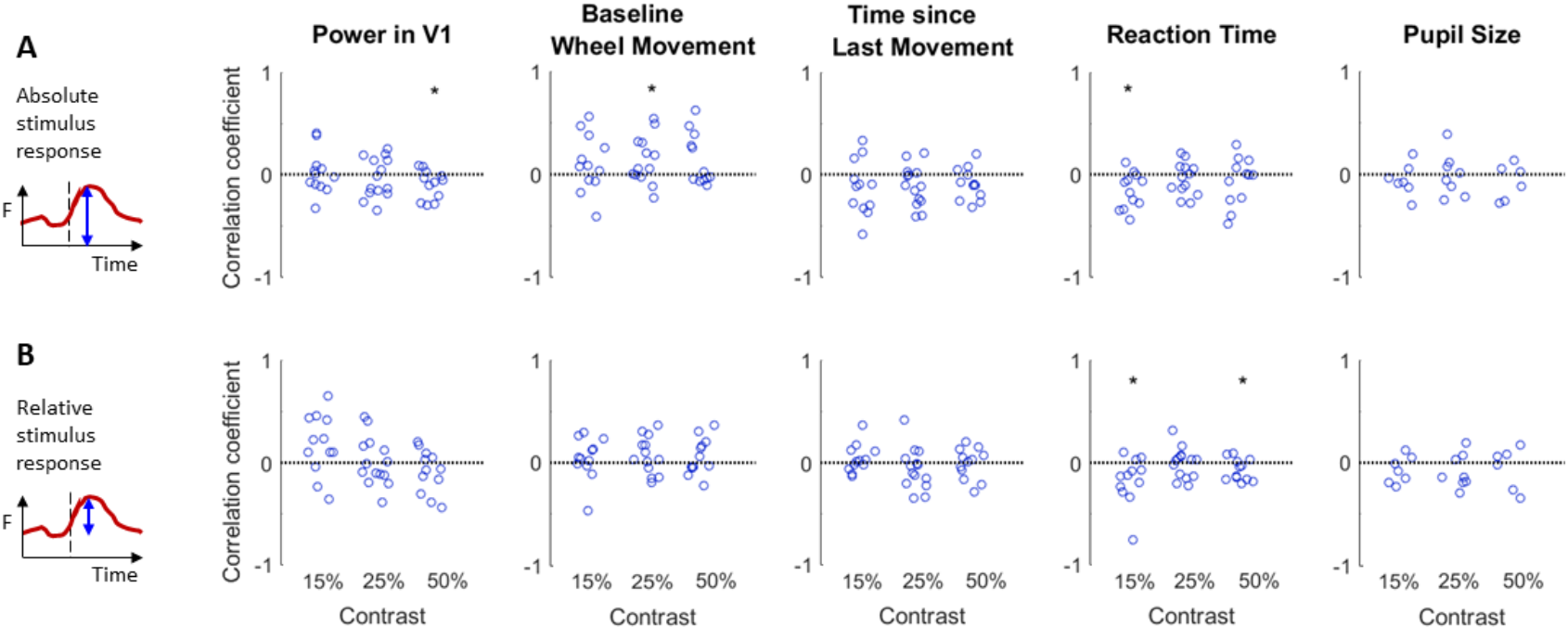
Stimulus response amplitude correlations with behavioral and physiological factors. Related to Figure 2. **A.** Absolute stimulus responses, with no baseline subtraction. **B.** Stimulus responses relative to baseline at time 0. Individual circles denote individual datasets.

**Supplementary Figure 3.**
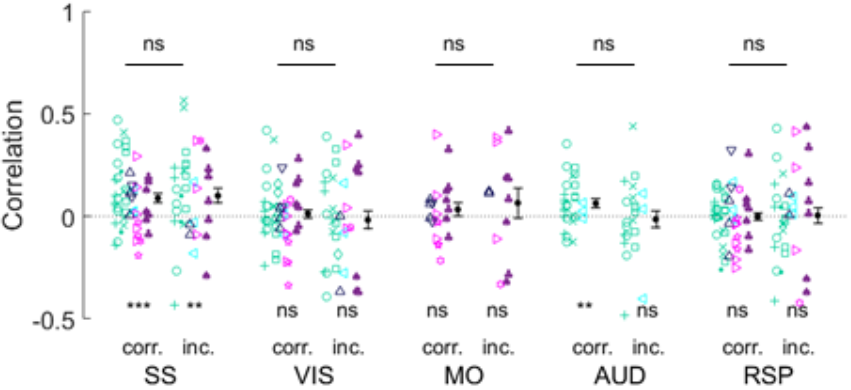
Correlation between 3-6Hz power and reaction time in correct (corr.; left) and incorrect (inc.; right) choices per ROI. Related to Figure 3. Color and symbol scheme is identical to the main figures (symbol color denotes genotype, glyphs denote individual animals). Black filled circles with bars indicate mean and SEM. *** = p<0.001, ** = p<0.01, * = p<0.05, ns = non significant.

**Supplementary Figure 4.**
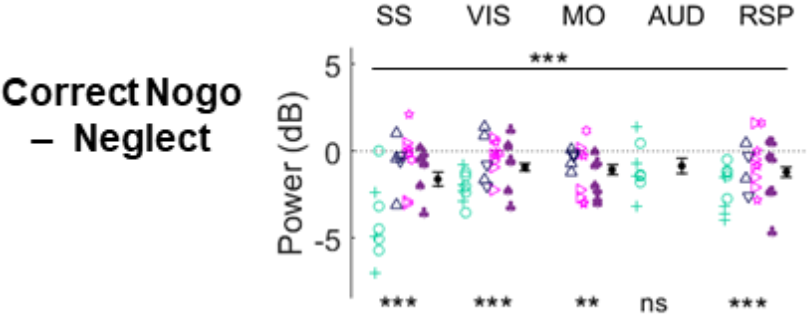
Summary of intercept differences between correct no-go and neglect trials. Related to Figure 4. Black filled circles with bars indicate mean and SEM. *** = p<0.001, ** = p<0.01, * = p<0.05, ns = non significant.

**Supplementary Figure 5.**
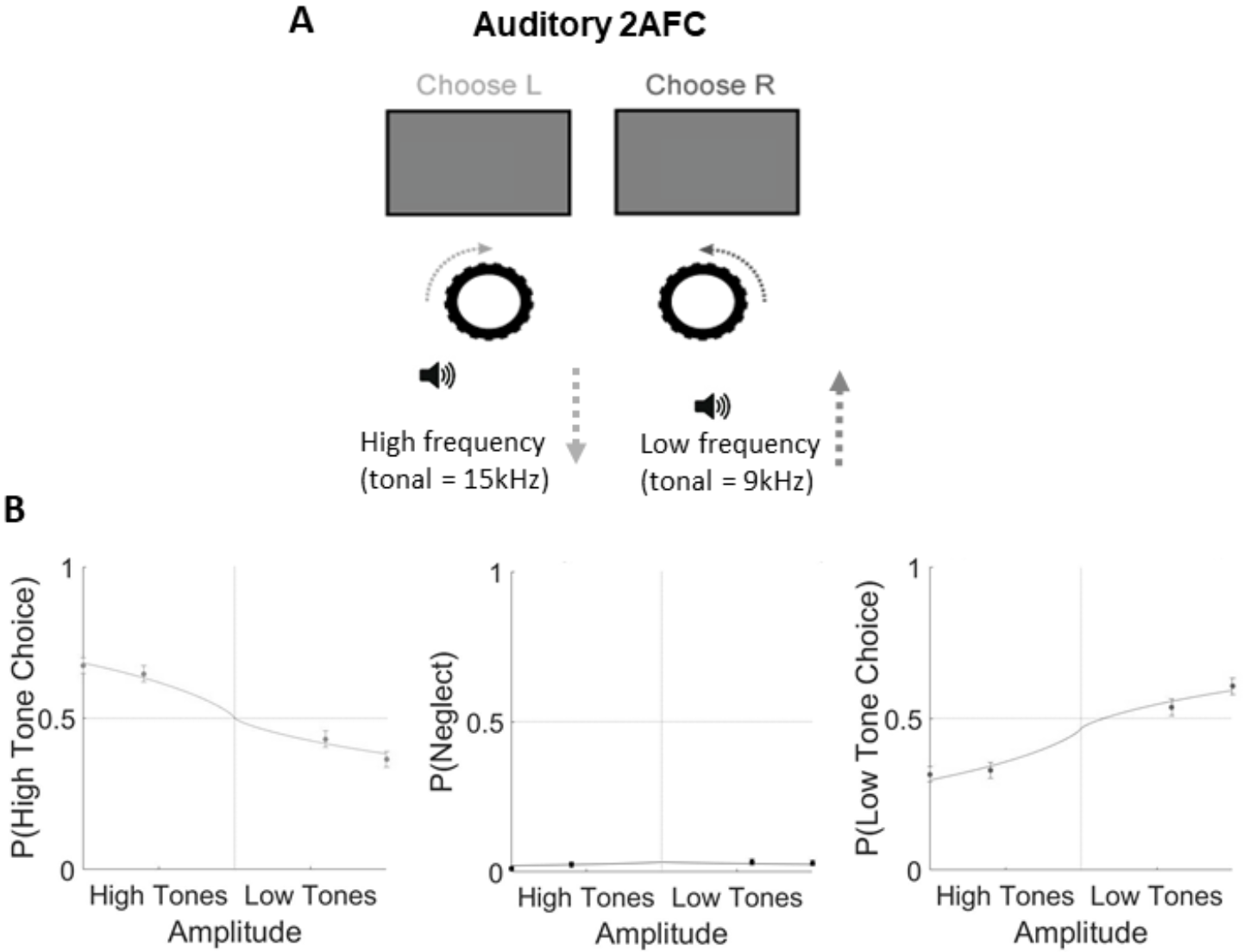
Auditory task performance. Related to Figure 7. **A.** Mice were headfixed in front of a grey iso-illuminant screen while auditory stimuli of either high or low tonal frequency were played from a speaker in front of them. Turning the wheel changed the tonal frequency of the stimulus, and the aim of the task was to bring the frequencies to a central target frequency. **B.** Example psychometric curves from one mouse. Filled circles and bars denote mean and SEM.

**Supplementary Figure 6.**
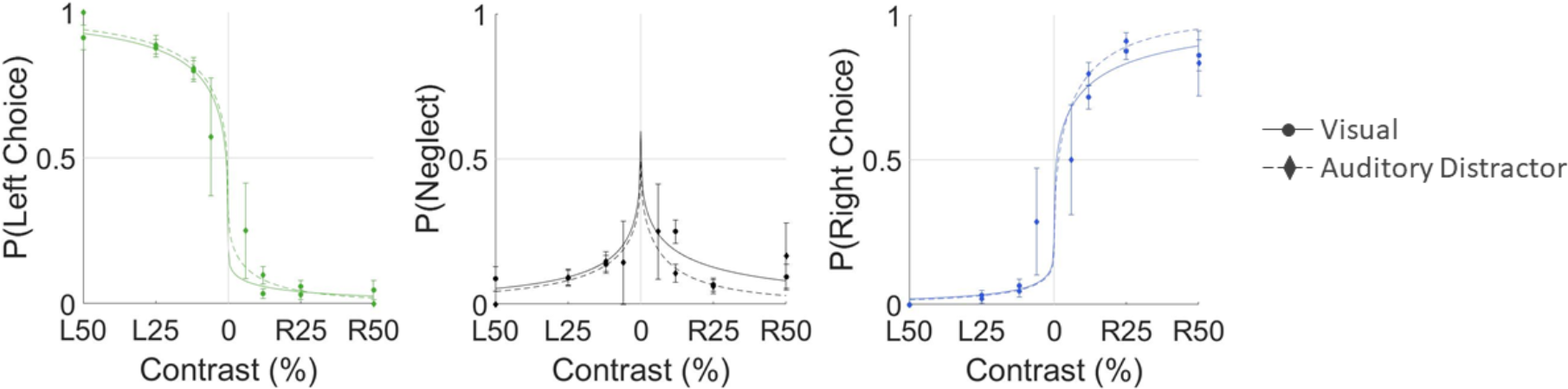
Psychometric curves comparing visual 2AFC and auditory distractor task performance. Related to Figure 7. Filled circles and bars denote mean and SEM.

**Supplementary Figure 7.**
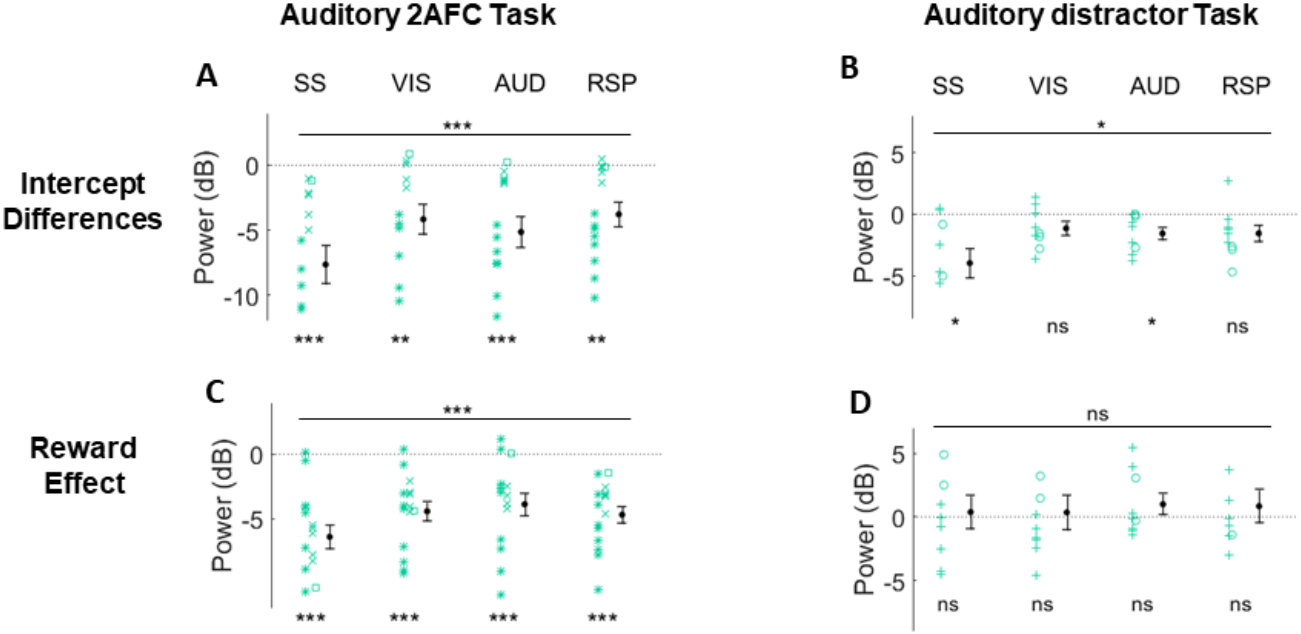
Summary intercept differences and reward effects across ROIs in the auditory 2AFC and auditory distractor tasks. Related to Figure 7. **A-B** Intercept differences, **C-D** reward effects. Black filled circles with bars indicate mean and SEM. *** = p<0.001, ** = p<0.01, * = p<0.05, ns = non significant.

**Supplementary Figure 8.**
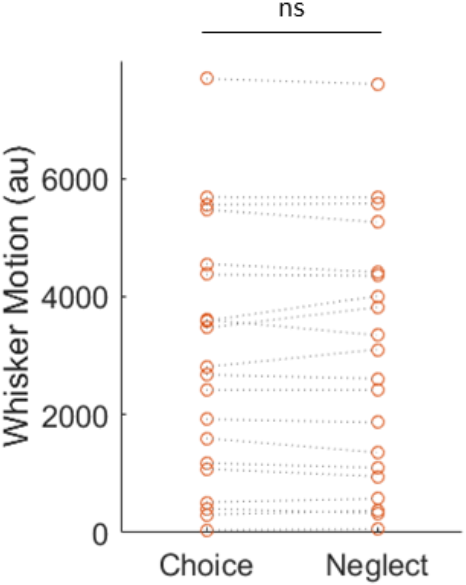
Quantification of whisker motion in choice and neglect trials. Whisker motion was quantified from motion energy in videos of one side of the face of the mouse. An ROI was defined close to the snout where the whiskers commence, and the whisker motion was computed as the absolute value of the difference of two frames within the ROI and summed over pixels greater than a threshold that was manually adjusted for each experiment. Each data point represents one experiment.

